# *Ex vivo* Immune Profiling in Patient Blood enables Quantification of innate immune Effector Functions

**DOI:** 10.1101/2020.10.01.322032

**Authors:** Teresa Lehnert, Ines Leonhardt, Sandra Timme, Daniel Thomas-Rüddel, Frank Bloos, Christoph Sponholz, Oliver Kurzai, Marc Thilo Figge, Kerstin Hünniger

**Author notes:** shared first authors contributed equally (Teresa Lehnert, Ines Leonhardt). shared senior authors contributed equally (Marc Thilo Figge, Kerstin Hünniger). **Correspondence:** Marc Thilo Figge, Kerstin Hünniger. **Author Contributions:** Conceptualization and Ideas: OK, MTF, KH; Formal Analysis including application of statistical, mathematical, computational, or other formal techniques: TL, ST, MTF; Funding Acquisition: OK, MTF; Investigation: TL, IL, ST, KH; Resources: DTR, FB, CS, Writing: TL, IL, OK, MTF, KH. **Funding:** This study was funded by the Federal Ministry for Education and Science (BMBF) within the Center for Sepsis Control and Care and by the Deutsche Forschungsgemeinschaft within the Collaborative Research Center Transregio 124 FungiNet “Pathogenic fungi and their human host: Networks of Interaction”, DFG project number 210879364 (projects B4 to MTF and C3 to OK).

## Abstract

The immune response towards infection is a dynamic system combating invading pathogens to maintain homeostasis and the integrity of the body. Unbalanced immune response profiles determine many clinical syndromes including sepsis and thus present a major challenge in management of life threatening infections. Consequently, there is a high demand to determine a patient’s immune status and identifying functional parameters for immune dysfunction.

Here, we quantified the global functional status of human innate immune responses by using a human whole-blood model of infection combined with biomathematical modeling. By determining functional parameters of innate immune cell populations after *ex vivo* whole-blood bacterial (*Staphylococcus aureus*) and fungal (*Candida albicans*) infection, we examined cell-specific functional parameters including migration rates or phagocytosis rates in patients that underwent cardiac surgery with extracorporeal circulation. This intervention is known to pose a transient but strong inflammatory stimulus. In addition to a post-operative increase in white blood cell count mainly caused by mobilization of immature neutrophils we find that the surgery induced pro-inflammatory stimulus results in a mitigation of pathogen-specific response patterns that are characteristic for healthy people and baseline results in our patients. Moreover, our model revealed changing rates for pathogen immune evasion, indicating increased inter-pathogenic differences after surgery. This effect was specific for *C. albicans* and could not be observed for *S. aureus*. In summary, our model gives insight into immune functionality and might serve as a functional immune assay to record and evaluate innate response patterns towards infection.

**Author summary:** Assessment of a patient’s immune function is critical in many clinical situations. One prominent example is sepsis, which results from a loss of immune homeostasis due to microbial infection. Sepsis is characterized by a plethora of pro- and anti-inflammatory simuli that may occur consecutively or simultaneously and thus any immunomodulatory therapy would require in depth knowledge of an individual patient’s immune status at a given time. Whereas lab-test based immune profiling often relies solely on quantification of cell numbers, we have used an *ex vivo* whole-blood infection model in combination with biomathematical modeling to quantify functional parameters of innate immune cells in patient blood. A small blood sample of patients undergoing cardiac surgery, which is known to constitute an inflammatory stimulus was infected *ex vivo*. Functional immune cell parameters were determined using a combination of experimental assays and biomathematical modeling. We show that these parameters change after an inflammatory insult triggered by cardiac surgery and extracorporeal circulation. This does not only interfere with pathogen elimination from blood but also selectively augments the escape of the fungal pathogen *Candida albicans* from phagocytosis.

## Introduction

Critical illness may be associated with significant alterations in immune function. Both hypo- and hyperinflammatory states are possible. While impairment of human immunity significantly increases the risk of infection, systemic inflammation promotes the development of the multiple organ dysfunction syndrome. Therefore, assessment of immune function would be desirable in clinical practice. However, only unspecific laboratory tests such as quantification of immune cell populations are available without providing information on immune cell function: Risk of infection during neutropenia is quantified by determining the number of neutrophils found in peripheral blood [1]. Similarly, the CD4^+^ T-cell count is used to quantify the degree of immunosuppression in HIV infection [2]. Although these assays are highly useful in clinical routine, they solely rely on quantitative thresholds and do not provide any information on immune cell function [3]. To improve understanding of a patient’s immune status, functional immune monitoring has been attempted by quantification of released proteins, such as IL-6, IL-8, procalcitonin and C-reactive protein [4]. However, these molecules are only indirect markers of cellular immunity. They derive from different immune cells, are mostly pleiotropic and may antagonize each other, all of which limits their clinical use. Another marker of immune dysfunction indicating sepsis-induced immunosuppression is the reduced expression of major histocompatibility (MHC) class II molecule HLA-DR on monocytes that is associated with a diminished antigen-presenting capacity and a shift from pro- to anti-inflammatory cytokine production (reviewed in [5,6]). Similarily, neutrophil surface expression of high-affinity Fcγ receptor I (CD64) has been shown to increase in patients during the early immune response to bacterial infection and in systemic inflammatory response syndrome [7– 9]. Several studies demonstrated that CD64 measured as an index may be useful for detection and management of sepsis and bacterial infection in neonatal intensive care units and adult hospital patients [10,11]. However, while surface markers reflect the activation status of immune cells, they do not provide a direct functional readout. Therefore, in a more complex setting, *ex vivo* stimulation of whole blood with LPS and quantification of cellular cytokine release has been used to quantify immune function. LPS-induced TNF-α release by monocytes in whole blood tends to be less in immunosuppressed patients compared to healthy individuals [12,13]. Nevertheless, this attempt so far only relied on a secreted cytokine as an indirect marker for cellular immune function. The most functional read-out described so far is the assessment of neutrophil dysfunction in critically ill patients by measuring neutrophil capacity to clear zymosan particles *ex vivo* [3,14]. However, these assays selectively address neutrophils and require prior isolation of the cells, which may alter their function especially in patients with activated neutrophils [15].

In previous studies, we applied a systems biology approach to investigate the immune response to pathogens in blood of healthy individuals [16–18]. Using *ex vivo* whole-blood infection in combination with biomathematical modeling enabled the calculation of functional parameters for blood immune cells, such as kinetic rates for pathogen uptake and killing. The aim of this study was to investigate wether a systems biology approach allows quantification of immune function in cardiac surgery patients after cardiopulmonary bypass. These patients receive a standardized anesthesia regiment and operation procedure and experience a well characterized inflammatory insult (e.g. increase of blood lipopolysaccharide and beta-D-glucan levels) which enables analysis of immune function before and after the insult [19–21].

## Results

### Pathogen-specific immune response patterns during bacterial and fungal whole-blood infection

Prior analyses in whole blood have exclusively been done with viable pathogens. However, quantification of immune cell function in patient blood samples requires the use of inactivated stimuli e.g. to avoid effects of pathogen inactivation by antibiotics in patient blood. Thus, we quantified immune cell functional parameters for two inactivated pathogens, *Staphylococcus aureus* (bacterial) and *Candida albicans* (fungal). During a 4-hour time course, neutrophils were the main immune cell type to interact with both pathogens (Fig 1A). Both pathogens showed similar association to neutrophils after 60 min of blood infection. However, the early 10 min time point revealed much faster association of *S. aureus* to neutrophils (70.8 ± 10.2%) when compared with *C. albicans* (45.0 ± 13.3%) and monocytes (7.6 ± 1.8% for *S. aureus* compared to 3.1 ± 0.5% for *C. albicans*, Fig 1A,B). Consistent with different association kinetics, *C. albicans* was cleared more slowly than *S. aureus* (10 min *p*.*i*.: extracellular *C. albicans* 52.9 ± 13.6%, extracellular *S. aureus* 21.6 ± 10.5%, Fig 1C). Notably, 4 hours after infection a larger population of *C. albicans* cells remained extracellular compared to *S. aureus* (extracellular *C. albicans* 13.0 ± 4.3%, extracellular *S. aureus* 4.0 ± 1.5%). Comparison of these data with previous studies and a new set of experiments using viable *C. albicans* showed overall comparable patterns but also revealed a significantly higher number of inactivated fungal cells associated to monocytes (60 min *p*.*i*.: viable *C. albicans* 4.9 ± 2.3%, inactivated *C. albicans* 13.4 ± 2.5%, *P* < 0.001) (Fig 1F and [16]). Association to lymphocytes could not be detected for either pathogen regardless of their viability. Whole-blood infection with *C. albicans* in an either active or inactive state induced a strong and comparable secretion of monocytes-derived cytokines (TNF-α, IL-1β and IL-6), whereas only low cytokine levels could be detected in mock-infected blood (Fig 1G).

**Fig 1.**
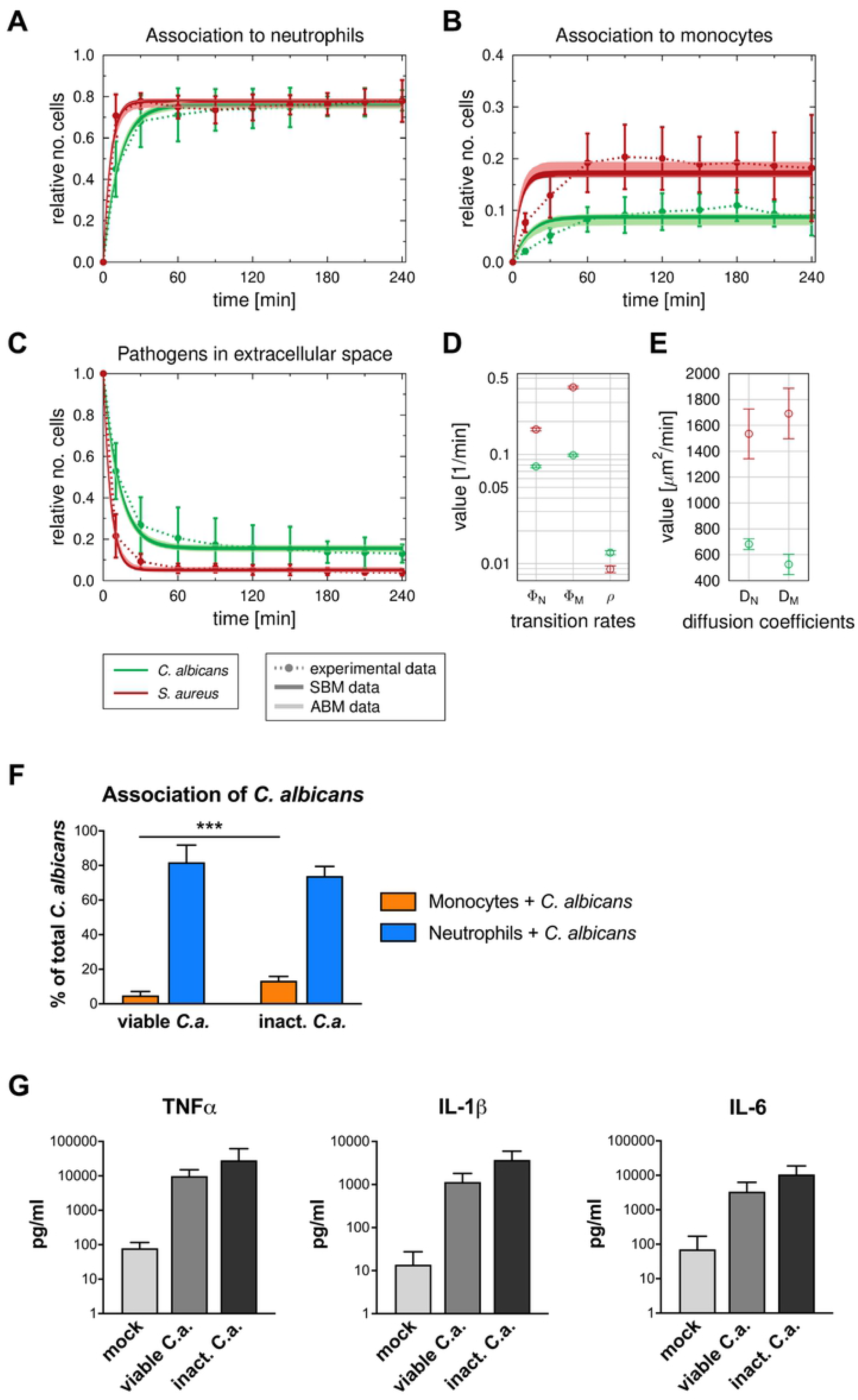
Comparison of the dynamics of host-pathogen interaction during *C. albicans* and *S. aureus* infection in healthy blood. (A-C) Results of fitting the state-based model (SBM) and the agent-based model (ABM) to experimental data of pathogen association to immune cells in whole-blood samples from healthy donors. The simulated dynamics of the combined units (solid lines) were obtained by fitting the SBM (dark color) and the ABM (light color) to the experimentally measured association kinetics (dotted lines). Experimental data were gained from whole-blood infection assays with either *C. albicans* (green lines) or *S. aureus* (red lines). Experimental data points and error bars refer to the means and standard deviations of 10 and 7 experiments, respectively, for *C. albicans* and *S. aureus*. The thickness of the solid lines showing the results from the SBM represents the standard deviations obtained by 50 simulations with transition rate values that were sampled within their corresponding standard deviation. The thickness of the solid lines showing the results from the ABM represents the standard deviations obtained from 30 stochastic simulations of the ABM with the estimated diffusion coefficients. (A) and (B), respectively, depict the dynamics of the combined units *P*_*N*_ and *P*_*M*_, which correspond to the experimental data on pathogen cells that are associated to neutrophils and monocytes. (C) shows the kinetics of the combined unit *P*_*E*_ together with experimentally measured kinetics of either fungal or bacterial cells that are in extracellular space. (D) Mean values (data points) and standard deviations (error bars) of transition rate values that were obtained by fitting the SBM to experimental data using the fitting algorithm simulated annealing. Here, the rate of phagocytosis by neutrophils (*ϕ*_*N*_) and by monocytes (*ϕ*_*M*_) as well as the rate for immune escape (*ρ*) are depicted for infection scenarios with either *C. albicans* (green data points) or *S. aureus* (red data points). (E) Diffusion coefficients for neutrophils (*D*_*N*_) and moncytes (*D*_*M*_) were estimated by fitting the ABM to the experimental data for both, *C*.*albicans* (green) and *S. aureus* (red), respectively. Mean and standard deviation are calculated from all parameter sets with a mean LSE that lies within the standard deviation of the optimal parameter set. (F) Association of viable and inactivated *C. albicans* with blood monocytes and blood neutrophils after 60 min of confrontation was quantified using flow cytometry. Values correspond to the means ± SD of at least five independent experiments with whole blood from different donors. (G) The release of monocyte-derived cytokines (TNF-a, IL-1β, IL-6) in plasma samples generated from 4 h whole-blood infection experiments in response to viable and inactivated *C. albicans* cells was investigated. Boxplots show means ± SD of at least three independent experiments with whole blood from different donors.

In order to comparatively quantify the functional properties of immune cells we calibrated the state-based virtual infection model (SBM) to the experimental data (see Methods section, S1 Table, Fig 1D). The coefficients of variation of the resulting transition rates are very small (< 7%) and the corresponding dynamics of the three combined units are similar to the mean values of the respective experimental data, indicating a good match to the experimental data (see Fig 1A-C). By comparing the resulting transition rate values, we observed significantly higher phagocytosis rates for *S. aureus* than for *C. albicans* (for neutrophils 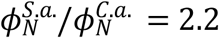 *P*′< 0.001, for monocytes 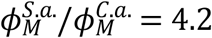 with *P*′< 0.001). The opposite ratio was found with respect to the rate of immune escape (*ρ*), (*ρ*^*S*.*a*.^/*ρ*^*C*.*a*.^ = 0.7 with *P′*< 0.001). These parameter differences lead to an overall faster removal of *S. aureus* cells from extracellular space and a higher amount of immune evasive *C. albicans* at 240 min after infection (Fig 1C). Based on the estimated parameters of the SBM we also simulated the infection scenario with both pathogens in the agent-based model. Similar to the SBM dynamics, the resulting dynamics of an agent-based virtual infection model (ABM) to the experimental data (see Methods section) are in agreement (see Fig 1A-C). Diffusion coefficients for neutrophils and monocytes are approximately two to three times higher for *S. aureus* infection (*D*_*N*_,*D*_*M*_) = (1500 μ*m*^2^/*min*,1800 μ*m*^2^/*min*) compared to *C. albicans* (*D*_*N*_,*D*_*M*_) = (700 μ*m*^2^ /*min*,450 μ*m*^2^/*min*). Furthermore, the diffusion coefficient for neutrophils is higher than for monocytes during *C. albicans* infection and *vice versa* for *S. aureus* infection (Fig 1E).

### Cardiac surgery results in neutrophil mobilization and immune activation

Data generated so far clearly show that a combination of *ex vivo* blood infection and virtual infection modelling allows quantification of immune cell functions. To address the feasibility of this approach in a clinical setting, we used blood samples from patients undergoing cardiac surgery for mitral valve insufficiency. Six patients (4 female, 2 male) with an age of 52 to 74 years were recruited. All received minimally invasive mitral valve replacement or reconstruction surgery via lateral thoracotomy on cardiopulmonary bypass (CPB). Anesthesia was induced by Propofol, Sufentanyl and Rocuronium and sustained by Sufentanyl and Sevofluran before and Propofol during CPB. Three patients underwent additional surgery such as tricuspid valve surgery, cryoablation, atrial appendage closure and atrial septal defect closure. Bypass time ranged from 99 to 220 minutes. All patients were extubated on the day of surgery and none had any major complications. The only postoperative infection was a urinary infection more than two weeks after surgery. After informed consent, blood samples were taken before cardiac surgery (pre-operative), directly after surgery (post-operative) and one day after admission to intensive care (post-operative + 1d). Both pro-inflammatory cytokines like IL-6, IL-8, MIP-1α and MIF as well as the anti-inflammatory cytokine IL-10 showed increased levels in post-operative blood (Fig 2A), whereas TNF-α and IL-1β were not detectable in plasma samples before and after surgery. Elevated levels were sustained until one day after surgery for IL-6 and IL-8. Furthermore, the mean total white blood cell counts increased after surgery (post-operative: 9.9 ± 0.8 × 10^9^/L, post-operative + 1d: 10.9 ± 1.0 × 10^9^/L) compared to pre-operative blood (4.7 ± 0.6 × 10^9^/L) (Fig 2B). Quantitative analyses of white blood cells revealed a significant increase of neutrophils (both post-operative and post-operative + 1d) and monocytes (post-operative + 1d), whereas lymphocyte counts remained stable. Neutrophilia following cardiac surgery is mediated by granulocyte colony-stimulating factor (G-CSF) induced bone marrow neutrophil release [22]. G-CSF was markedly higher in plasma samples obtained directly and one day after surgery compared to pre-operative blood (Fig 2A). Furthermore, whereas pre-operative blood was characterized by a homogenous neutrophil population that expressed abundant CD10 and high CD16 surface levels (mature neutrophils), the neutrophil population after surgery contained a CD10-negative subpopulation with reduced CD16 and increased L-selectin (CD62L) expression, indicating recruitment of immature neutrophils (S1A Fig). Post-operatively, the CD10^neg^ neutrophil subpopulation accounted for 49.3 ± 4.1% of the total neutrophil population and remained almost stable after one day (46.7 ± 2.6%). Whereas no changes in monocyte counts could be observed between the pre- and post-operative time point, total monocyte numbers were significantly increased one day after surgery (pre-operative: 0.24 ± 0.06 × 10^9^/L, post-operative + 1d: 0.54 ± 0.09 × 10^9^/L, *P* < 0.05, Fig 2B). Expression of the MHC class II antigen HLA-DR on monocytes was markedly decreased after surgery (S1B Fig). Together with the increased CD62L expression this pointed towards an immature state of monocytes induced by the surgery.

**Fig 2.**
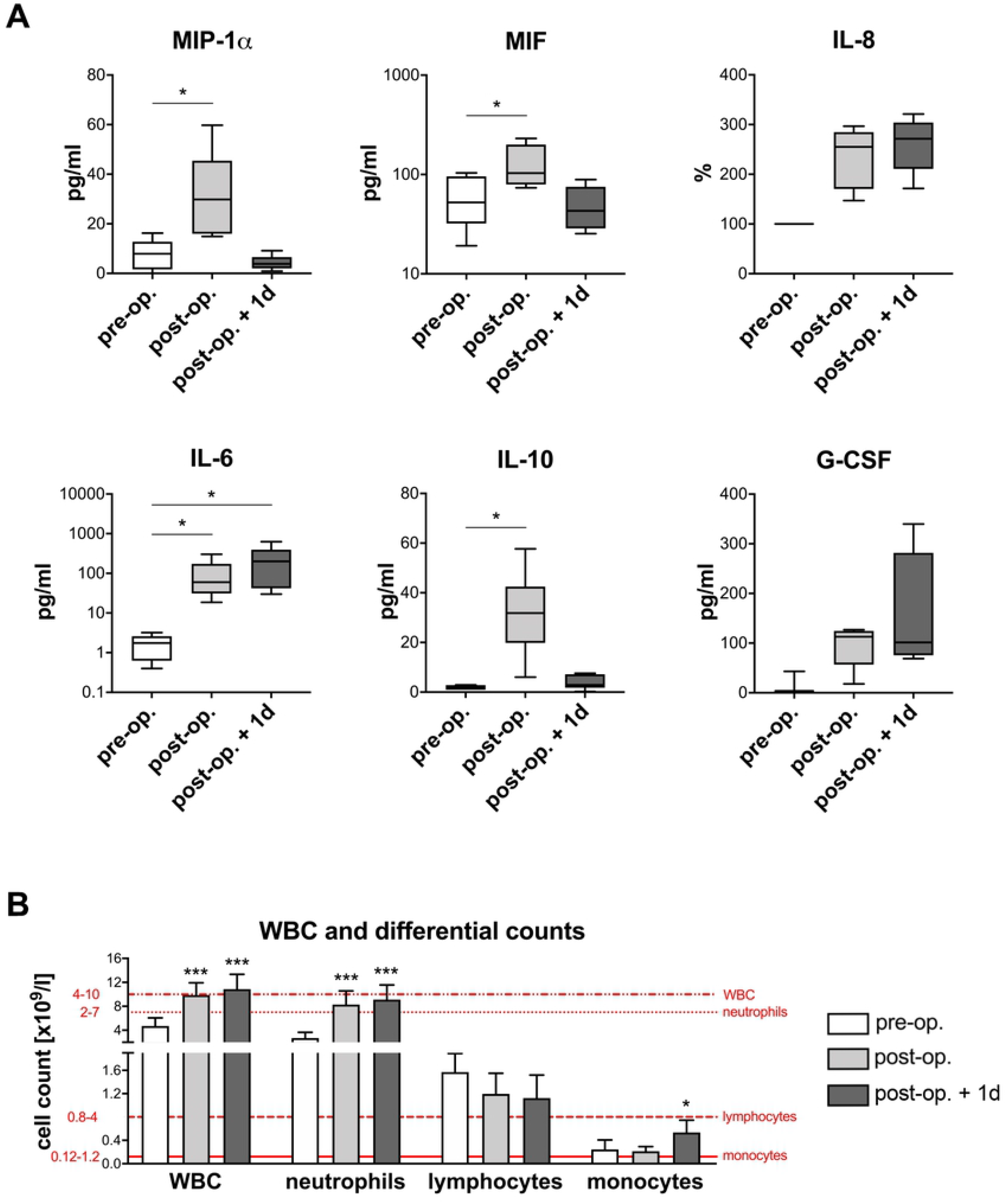
Blood after surgery shows changes in cytokine profiles and peripheral differential cell counts. Blood samples from six HLM patients taken before cardiac surgery (pre-operative), immediately after surgery (post-operative) and one day after admission to intensive care (post-operative + 1d) were analyzed for (A) cytokines levels (MIP1α, MIF, IL-8, IL-6, IL-10, G-CSF) using Luminex technology and (B) white blood cell count (WBC) as well as neutrophil, lymphocyte and monocyte counts using a automated hematology analyzer. Reference ranges of leukocytes are indicated in red. Significance was estimated using the unpaired, two-sided Student *t* test and shown as \**P* < 0.05, \*\*\**P* < 0.001.

### Pro-inflammatory stimulus during cardiopulmonary bypass mitigates inter-patient and inter-pathogen differences in immune response patterns

To quantify functional immune parameters in whole blood after cardiopulmonary bypass, we perfomed whole-blood infection with inactivated *S. aureus* or *C. albicans* as described before. These analyses revealed clear changes in the patterns of the innate immune response to *C. albicans* and *S. aureus*. After surgery, pathogen association to neutrophils and monocytes was faster and higher in comparison to the pre-operative time point (Fig 3). After 10 min post infection 35.1 ± 8.8% of *C. albicans* and 53.7 ± 7.2% of *S. aureus* cells, respectively, were associated to neutrophils in pre-operative blood (Fig 3A,D), whereas these fractions increased to 78.5 ± 5.1% for *C. albicans* and 81.1 ± 6.7% for *S. aureus* one day after surgery (Fig 3C,F). To test, whether altered association kinetics could be explained by altered leukocyte numbers or indicate functional alterations, we fitted the SBM to the association kinetics and used the means of the measured immune cell counts (Table S2) as input for the model simulations. These analyses clearly showed that the faster association kinetics after surgery can only be explained by a significant increase in phagocytosis rates of neutrophils and monocytes and cannot solely be explained by the increase of immune cell counts (Fig 3G). Importantly, pathogen-specific response patterns that were clearly visible for healthy donors and in samples taken before surgery were assimilated after the pro-inflammatory stimulus. More specifically, *S. aureus* had a greater magnitude and faster association to neutrophils and monocytes than *C. albicans* in blood taken before surgery (Fig 3A,D). Comparing the transition rates for phagocytosis by neutrophils (*ϕ*_*N*_) and monocytes (*ϕ*_*M*_) between both pathogens revealed significantly higher values for *S. aureus* 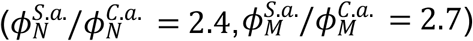 (Fig 3G). In contrast, association kinetics for *C. albicans* and *S. aureus* as well as the corresponding phagocytosis rates for neutrophils and monocytes were almost similar in blood after surgery (e.g. post-operative + 1d: 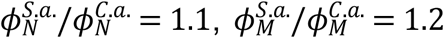; Fig 3G).

**Fig 3.**
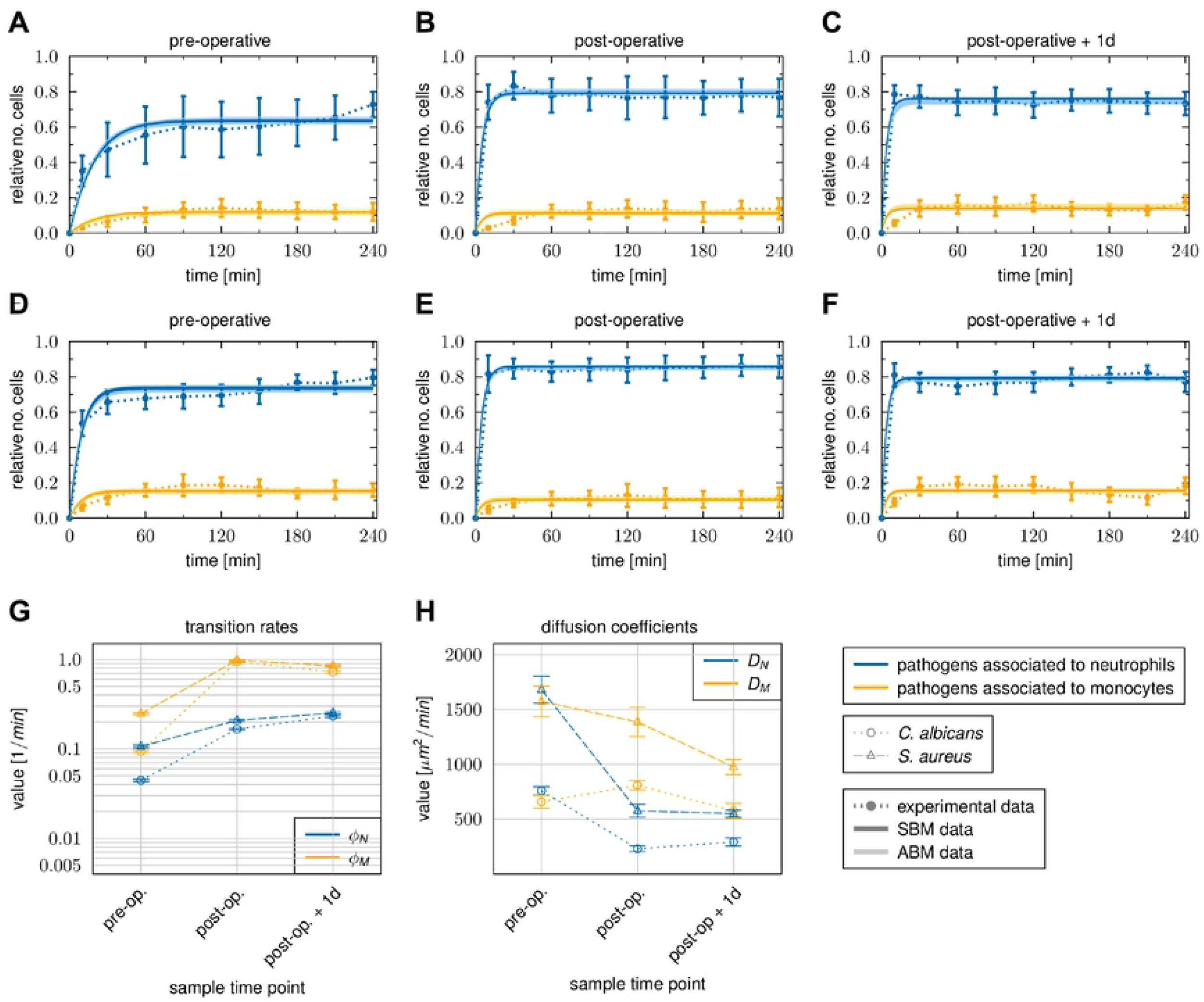
Time courses of pathogen association to immune cells observed in whole-blood samples of HLM patients. Blood samples were taken before cardiac surgery (pre-operative), immediately after surgery (post-operative) and one day after admission to intensive care (post-operative + 1d). Time-resolved experimental data (dotted lines) were obtained by whole-blood infection assays with either *C. albicans* (A-C) or *S. aureus* (D-F). Data points and error bars refer to the means and standard deviations of blood samples from six HLM patients. The simulated dynamics of the combined units (solid lines) were obtained by fitting the state-based model (SBM, dark color) and the agent-based model (ABM, light color) to the experimental data. The thickness of the results from the SBM (solid lines, dark color) represents the standard deviations obtained by 50 simulations with transition rate values that were sampled within their corresponding standard deviation. The thickness of the results from the ABM (solid lines, light color) represents the standard deviations obtained from 30 stochastic simulations of the ABM with the estimated diffusion coefficients. (G) Transition rate values of the SBM resulting from fitting the model to experimental data of either *C. albicans* or *S. aureus* infection in blood samples from HLM patients. The transition rate values are given for the phagocytosis rate *ϕ*_*N*_ of neutrophils and the phagocytosis rate *ϕ*_*M*_ of monocyte. (H) The diffusion coefficients are given for neutrophils *D*_*N*_ and monocytes *D*_*M*_. Mean and standard deviation are calculated from all parameter sets with a mean LSE that lies within the standard deviation of the optimal parameter set.

We also investigated possible alterations in immune cell migration in blood after surgery compared to the pre-operative time point using the ABM. As in healthy donors, diffusion coefficients for both immune cell types during *S. aureus* infection exceed diffusion coefficients during *C. albicans* infection by a factor of approximately two to three (Fig 3H, S2 and S3 Figs). In post-operative blood we observed for *C. albicans* infection a decrease in *D*_*N*_ by a factor of 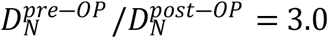 and an increase in *D*_*M*_ by a factor of 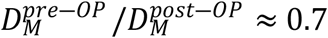 (Fig 3H and S2 Fig). Similarly, we also observed a decrease in *D*_*N*_ during *S. aureus* infection by a factor of 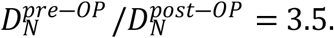 Contrary to *C. albicans* infection we did not observe an increase in *D*_*M*_ in post-operative blood, but found a slight decrease in *D*_*M*_ (Fig 3H and S3 Fig). Furthermore, we found that for both species *D*_*N*_ did not change from post-operative time point to one day after surgery. However, in case of *D*_*M*_ we observed a decrease for both species by a factor of approximately 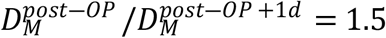. Comparing the values for *D*_*N*_ and *D*_*M*_ between the pathogens for each sample time point showed that both diffusion constants are larger for *S. aureus* infection in comparison to *C. albicans* infection for all time points. However, similar to the immune cell association and phagocytosis rates, the differences between the pathogens decrease after surgery for the diffusion of neutrophils and monocytes (see Fig 3H). In addition to assimilation of differences between both pathogens, the inter-individual differences in the association to immune cells decreased after the inflammatory stimulus. For instance, the percentage standard deviation of pathogens that interacted with neutrophils decreased from 32% to 10% for *C. albicans* and 13.5% to 8% for *S. aureus* infection between pre-operative and one day after surgery samples.

Taken together, kinetics of immune responses are accelerated after the pro-inflammatory stimulus independent of the microbial trigger. This results not only from the large increase in immune cell numbers, but also from increased transition rates (see Fig 3G). As a consequence pathogen specific -as well as the inter-individual differences are less pronounced after surgery.

### Monocyte activation by both *C. albicans* and *S. aureus* is dampened after cardiopulmonary bypass

We further investigated consequences of cardiopulmonary bypass surgery-induced inflammation on activation of innate immune cells during infection. Activation of monocytes analyzed by their surface expression of early activation antigen CD69 and cytokine secretion in response to microbial confrontation was reduced after cardiac surgery. Presence of *C. albicans* and *S. aureus*, respectively, induced the release of monocytic cytokines TNF-α, IL-1β and IL-6 within blood from all three time points tested for HLM patients compared to the corresponding mock-treated samples (Fig 4A). However, the induced levels were markedly lower in post-operative blood during both *C. albicans* (e.g. TNF-α: pre-operative 1456 ± 1248 pg/ml, post-operative 355 ± 428 pg/ml, *P =* 0.068) and *S. aureus* infection (e.g. TNF-α: pre-operative 1730 ± 903 pg/ml, post-operative 380 ± 427 pg/ml, *P* < 0.01). The reduced pro-inflammatory cytokine release pointed towards a regulatory effect mediated by increased IL-10 levels, which could be detected specifically in blood obtained directly after surgery (see Fig 2A), on monocytes. Neither *C. albicans* nor *S. aureus* infection were able to further induce IL-10 secretion when compared to mock-treated post-operative samples (Fig 4A). Since plasma concentrations of IL-10 almost returned to values within pre-operative blood after one day at the ICU, release of monocytic cytokines was also increased again in response to both pathogens (e.g. TNF-α: *C. albicans* 1229 ± 1234 pg/ml, *S. aureus* 1507 ± 852 pg/ml). In line with cytokine data, elevated CD69 surface levels on monocytes could be detected 4 hours after inoculation of *C. albicans* (MFI 1276 ± 598) or *S. aureus* (MFI 1447 ± 597) compared to mock-infection (MFI 488 ± 298) in pre-operative blood without any quantitative differences between both pathogens (Fig 4B). A less pronounced response was detected in blood after surgery (e.g. *S. aureus* infection: post-operative MFI 524 ± 202, post-operative + 1d MFI 580 ± 212). Together with reduced monocytic cytokine release, these data indicated a lower monocyte stimulation following infection of blood after surgery. Analyses of the surface phenotype of neutrophils revealed significantly differences in exposure of CD16 and CD66b between mock-infected control samples in blood taken before and after surgery as well as after one day at the ICU that can be explained by the inflammation-dependent recruitment of immature neutrophils (Fig 4B). Activation was largely restricted to those neutrophils that had phagocytosed either *C. albicans* or *S. aureus* and was stronger in response to *C. albicans*, shown by a more pronounced increase for CD69 (pre-operative: mock-infected MFI 141 ± 82, *C. albicans*-infected MFI 730 ± 178, *S. aureus*-infected MFI 427 ± 211) and down-regulation of CD16 (pre-operative: mock-infected MFI 7359 ± 2962, *C. albicans*-infected MFI 1721 ± 998, *S. aureus*-infected MFI 3812 ± 2108). Despite different levels in the control samples, the maximum of CD16 decrease and degranulation marker CD66b up-regulation on pathogen-associated neutrophils, respectively, was equal within blood from all three time points. Comparable results could be observed for CD69 expression.

**Fig 4.**
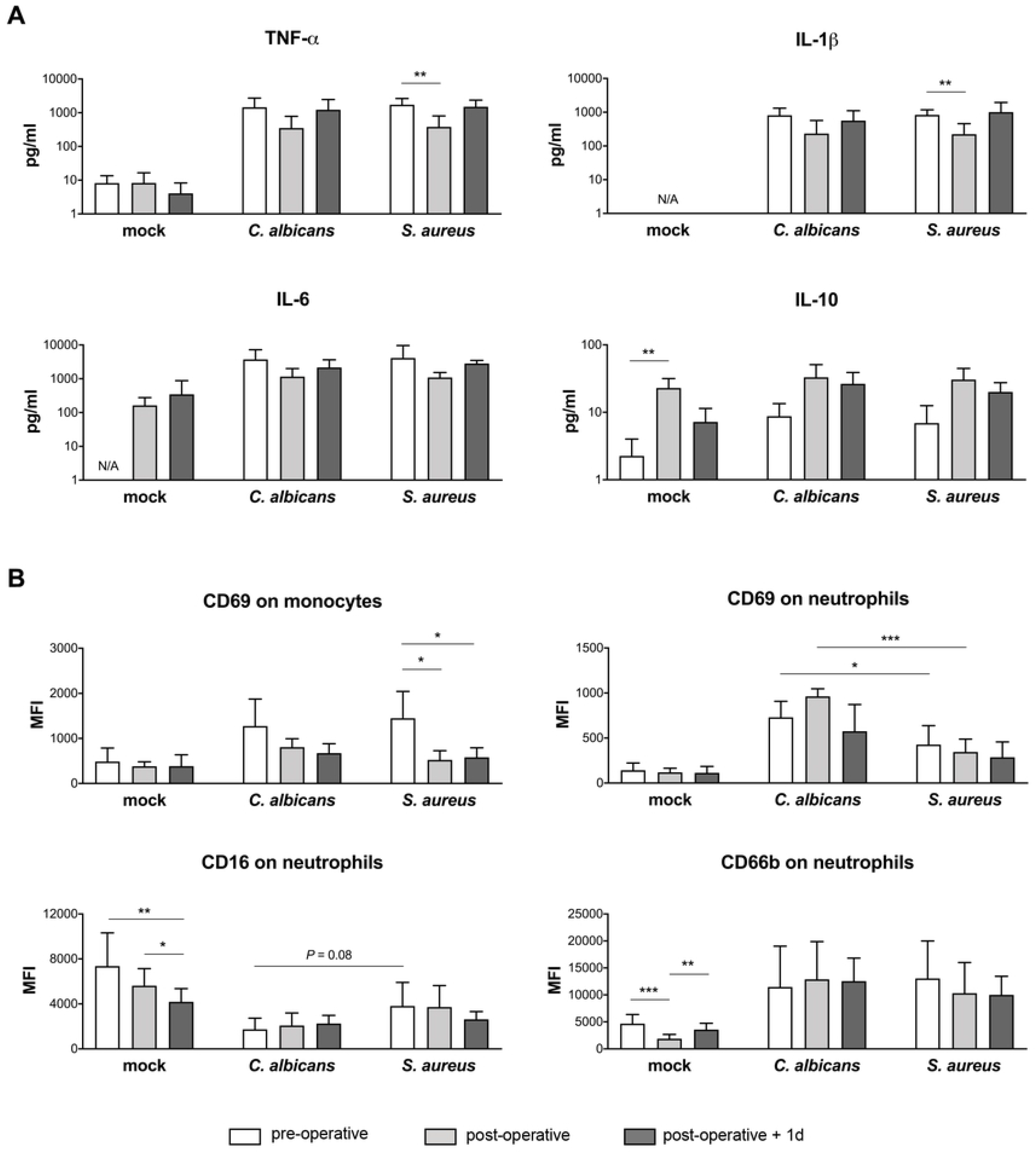
Changes in cytokine secretion and innate immune cell activation in whole blood from HLM patients after surgery. Blood samples from HLM patients were taken before cardiac surgery (pre-operative, non-filled bars), directly after surgery (post-operative, light grey bars) and one day after admission to intensive care (post-operative + 1d, dark grey bars) and either mock-infected, treated with *C. albicans* or *S. aureus* for 4 hours. (A) Plasma levels of TNF-α, IL-1β, IL-6 and IL-10 were quantified and bars are shown as means ± SD. Results are presented as pg/ml; N/A stands for values not available. (B) Surface marker expression was analyzed on total monocyte population and on pathogen-associated neutrophils by flow cytometry. Data shown are mean fluorescence intensity (MFI) ± SD. Significance is shown as \**P* < 0.05; \*\**P* < 0.01; \*\*\**P* < 0.001, unpaired, two-sided Student *t* test.

### Inflammatory stimulus during cardiopulmonary bypass induces enhanced *C. albicans* immune escape

Despite the quantitative increase in leukocyte number and functional changes, a fraction of *C. albicans* and *S. aureus* cells, respectively, was not phagocytosed by neutrophils or monocytes even after 4 hours of infection and remained extracellular (Fig 5). Comparing the fraction of extracellular cells between the different sample time points, we observed for both pathogens that this population decreased after surgery and the slope of the reaction curve during initial phase is increased. More precisely, 60 min after inoculation 34.5 ± 17.5 % of total *C. albicans* cells were not associated to immune cells in pre-operative blood (Fig 5A). However, in blood samples from immediately (post-operative) and one day after surgery only 9.9 ± 7.8% and 10.1 ± 3.7 % *C. albicans* cells, respectively, were not associated to immune cells at 60 min post infection (Fig 5B,C). Although these results indicated that elimination of pathogens from extracellular space after surgery was more efficient, pathogen-specific differences were still present within blood of the three tested conditions: in every case the fraction of extracellular cells was higher for *C. albicans* than for *S. aureus* following 240 min of infection (Fig 5A-C). Our model predicted that at 240 min all extracellular cells for both pathogens acquired immune escape (*P*_*IE*_). Interestingly, despite increased neutrophil numbers and neutrophil phagocytosis rates in blood after surgery, estimated transition rates for acquiring immune escape (*ρ*) were 2-fold higher for *C. albicans* in post-than pre-operative samples, which was in contrast to *S. aureus* with almost equal *ρ* values for the different sample time points (Fig 5D, S3 and S4 Tables). In fact, this resulted in a changed ratio of *ρ* between *S. aureus* and *C. albicans* infection in post-operative blood. While *ρ* values were similar for *C. albicans* and *S. aureus* in pre-operative blood (*ρ*^*S*.*a*.^/*ρ*^*C*.*a*.^ = 0.96), larger differences could be detected in blood samples taken after surgery (post-operative: *ρ*^*S*.*a*.^/*ρ*^*C*.*a*.^ = 0.48, post-operative + 1d: *ρ*^*S*.*a*.^/*ρ*^*C*.*a*.^ = 0.5).

**Fig 5.**
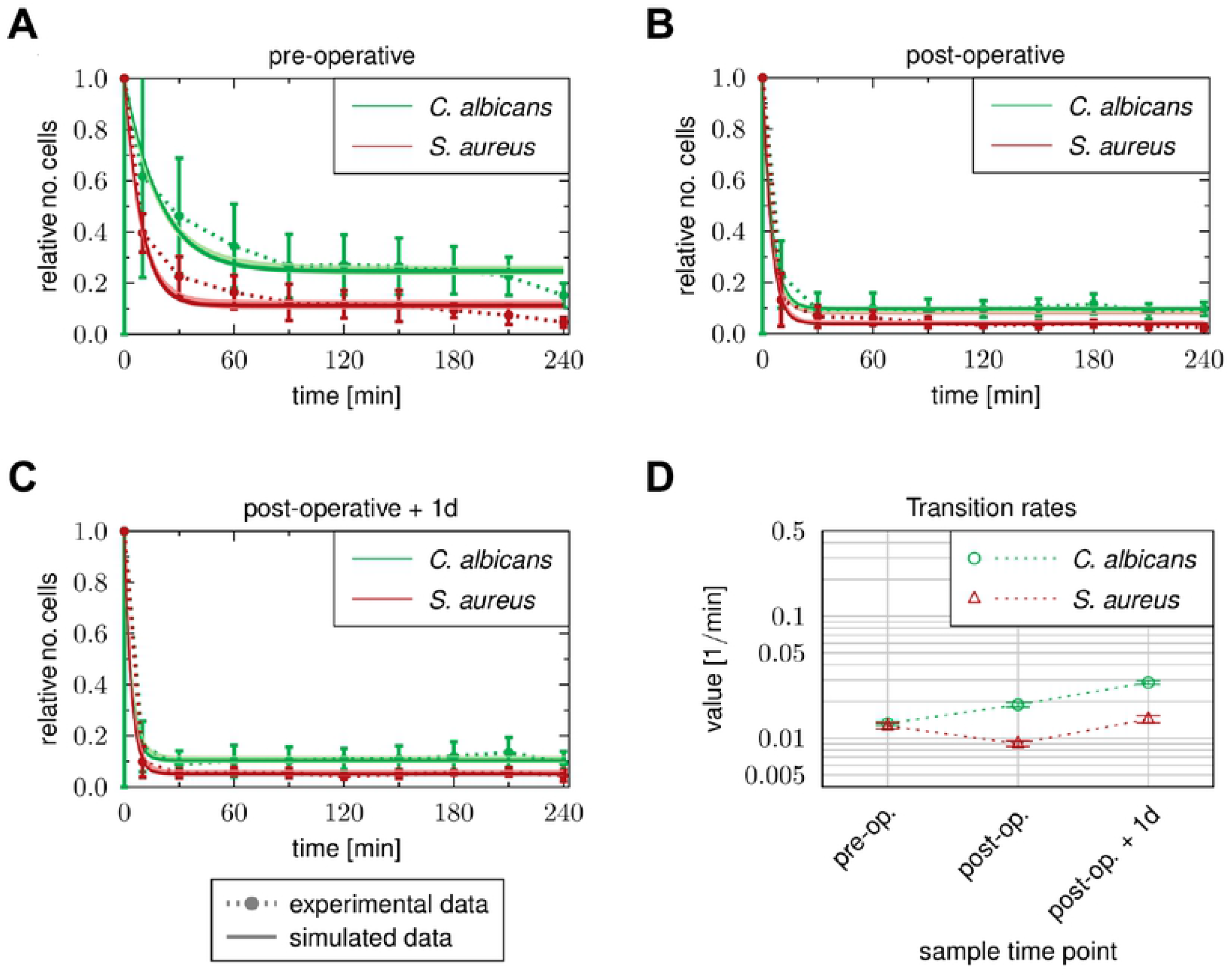
Time courses of extracellular pathogen association to immune cells observed in whole-blood samples of HLM patients. Blood samples were taken (A) before cardiac surgery (pre-operative), (B) immediately after surgery (post-operative) and (C) one day after admission to intensive care (post-operative + 1d). Time-resolved experimental data (dotted lines) were obtained by whole-blood infection assays with either *C. albicans* (green) or *S. aureus* (red). Data points and error bars refer to the means and standard deviations of blood samples from six HLM patients. The simulated dynamics of the combined units (solid lines) were obtained by fitting the state-based model (SBM, dark color) and the agent-based model (ABM, light color) to the experimental data. The thickness of the results from the SBM (solid lines, dark color) represents the standard deviations obtained by 50 simulations with transition rate values that were sampled within their corresponding standard deviation. The thickness of the results from the ABM (solid lines, light color) represents the standard deviations obtained from 30 stochastic simulations of the ABM with the estimated diffusion coefficients. (D) Transition rate values for immune evasion *ρ* of the SBM resulting from fitting the model to experimental data of either *C. albicans* or *S. aureus* infection in blood samples from HLM patient.

## Discussion

In this study, we applied a previously established systems biology approach, where experimental whole-blood infection assays are combined with virtual infection modeling to investigate the effect of immune activation on the immune defense against pathogens in whole blood. In order to analyze the immune response of patients that suffer from an unbalanced immune homeostasis, we used blood samples of patients that underwent cardiac surgery with extracorporeal circulation. The surgical intervention and the cardiopulmonary bypass are known to trigger a systemic inflammatory response. Whereas this transitory inflammation is not usually related to infection or sepsis, it can contribute to post-operative complications such as organ dysfunction or bleeding [23]. Pathophysiologically, a multitude of triggers that include surgical trauma, ischemia and reperfusion injury as well as endotoxinemia and glucanemia results in an acute phase reaction largely but not exclusively mediated by activation of NF-κB [20,24]. In particular, the inflammatory stimulus causes a surplus of the complement protein C5a, which serves as an important chemoattractant of several innate immune cells like neutrophils and monocytes [25,26]. C5a molecules activate neutrophils and induce neutrophil degranulation and superoxide generation [26–32]. Due to this pro-inflammatory stimulus, patients after cardiac surgery show leukocytosis, including mobilization of neutrophils from bone marrow. Furthermore, their monocytes express reduced levels of HLA-DR and increased CD62L [33]. Thus, transient hyperinflammation can well be studied in elective cardiac bypass surgery patients.

In accordance with this, we could observe leukocyte recruitment after surgery in our study. Especially the number of neutrophils increased by more than twice and was due to recruitment of CD10^−^ immature neutrophils. In addition, monocytes were increased at day one, but showed an immature phenotype already in blood taken directly after surgery. Despite the higher abundance of immature neutrophils and monocytes, pathogen association to immune cells occured faster and to a larger extend after then before surgery. By calibrating the state-based virtual infection model to the association kinetics of samples taken before and after surgery, we found that this was not solely due to increased immune cell numbers. In contrast, phagocytosis rates of neutrophils and monocytes increased by a factor of more than two after surgery, indicating an enhanced activation status. These quantitative results showing the increase of immune cell activity could not be detected by selectively quantifying surface activation marker expression, such as for the neutrophil degranulation marker CD66b. Nevertheless, despite the high abundance of immature neutrophils recruited in response to transient hyperinflammation, these cells showed the same maximum of CD66b and CD69 upregulation and decrease in surface CD16 upon fungal phagocytosis compared to mature neutrophils in pre-operative blood. Immature neutrophils already contain all three types of granules and can produce reactive oxygen species [34,35], indicating that these cells are a functional subset that becomes released in certain conditions. Studies by Leliefeld *et al*. and van Grinsven *et al*. using experimentally induced acute inflammation in healthy subjects by intravenous administration of bacterial LPS demonstrated that the different neutrophil subsets not only differ in phenotype but also in function [36,37]. Immature neutrophils were shown to have even higher antibacterial capacity *in vitro* and exhibit efficient migration, which led to the hypothesis that immature neutrophils are not released as bystanders but rather as cells more efficient in pathogen killing, in line with the observed increased immune cell activity in our study. Other previous studies reported an increase of immune cell activation for monocytes and neutrophils after surgery, by evaluating the expression of immune cell specific activation markers and the cytokine profile [38–40]. However, they do not provide a quantitative change of the immune cell functionality during the response to different pathogens serving as stimuli. The observed functional activation also mitigates pathogen specific patterns of immune activation: Prior to the inflammatory insult and as observed in samples from healthy control donors, *S. aureus* infection induced a faster immune response than *C. albicans* infection. In contrast, after surgery, the inter-pathogen differences were substantially reduced, indicating that immune responses in blood after inflammatory insult show less specificity for the trigger pathogen.

Additionally, we analyzed the concentration of pro- and the anti-inflammatory cytokines in blood samples of HLM patients before and after surgery. Interestingly, pro-inflammatory cytokines IL-1β and TNF-α were not present in higher levels after surgery, whereas an increase was observed for others, such as IL-6. In addition, the anti-inflammatory cytokine IL-10 was also significantly increased at the end of surgery, indicating the activation of anti-inflammatory pathways. These findings are in line with several studies reporting the surgery-induced release of both pro- and anti-inflammatory mediators [27,41–44]. During *ex vivo* infection, the release of monocytic cytokines as well as up-regulation of early activation antigen CD69 on monocytes were markedly lower in post-operative blood. Together with the immature phenotype of blood monocytes this pointed towards a regulatory effect of increased IL-10 levels in blood taken directly after surgery. Among the mediators released during surgery, IL-10 was already shown to contribute to the down-regulation of HLA-DR on CD14^+^ cells [45]. IL-10 levels decreased after one day, which consequently resulted in the partial recovery of HLA-DR on monocytes and secretion of cytokines IL-1β, IL-6 and TNF-α during both *C. albicans* and *S. aureus* infection.

Similar to results of our previous studies on the immune response in whole-blood samples upon *C. albicans* infection [16], we observed a specific population of pathogens that could not be phagocytosed by the immune cells, but are still present in extracellular space at four hours after infection. Therefore, these pathogens can escape the immune defense. Even though, the evasive pathogens could be detected in stimulated blood samples at each sample time point before and after surgery, we found that the amount of evasive *C. albicans* cells and *S. aureus* cells decreased after surgery, while the rate for this process increased for *C. albicans* infection and remained almost unchanged for *S. aureus* infection. In a previous study, we investigated the mechanism of *C. albicans* immune evasion in whole blood by testing potential evasion mechanisms using mathematical modeling. By simulating the infection in whole blood under neutropenic conditions, we could suggest future experimental measurements that most likely enable to accept or reject one of the two tested mechanisms, *i*.*e*. spontaneous evasion or neutrophil-mediated immune evasion. Since we observed larger evasion rates after surgery, where the immune cells are more active in terms of phagocytosis, our study provides clues that the evasion of pathogens could be dependent on immune cell activity. Moreover, the change in the immune evasion rate after surgical insult is an additional characteristic property describing the changes of the immune response in whole-blood samples of patients that underwent inflammatory stimulus. Taken together we show that a combination of experimental assays in *ex vivo* blood samples and biomathematical monitoring is a feasible approach for immune profiling in patients. Clearly at this stage this cannot be used for diagnostic purposes and requires both further optimization and standardization. However, using this approach in its current state we can already reveal important pathophysiological properties governing virulence of microorganisms as in the case of immune escape of *Candida albicans*.

## Material and Methods

### Experimental methods

#### Patients and Ethics statement

Human peripheral blood was collected from healthy volunteers and HLM patients with written informed consent. This study was conducted in accordance with the Declaration of Helsinki and all protocols were approved by the Ethics Committee of the University Hospital Jena (permit number: 4643-12/15). Patients included in this study received standardized minimally invasive cardiac surgery for mitral valve insufficiency with a heart lung machine (HLM) and same anesthesia regiment. Blood was taken from inserted catheters before cardiac surgery (pre-operative), immediately after surgery (post-operative) and one day after admission to the intensive care unit (post-operative + 1d).

#### Strains and Culture

*Candida albicans* (SC5314) was grown overnight in YPD medium (2% D-glucose, 1% peptone, 0.5% yeast extract, in water) at 30 °C to stationary phase. Fungal cells were reseeded in YPD medium, grown for 3 hours at 30 °C, stained with CellTracker^™^ Green (Invitrogen) for 1 hour, and harvested in PBS. Subsequently, cells were inactivated with 0.1% thimerosal (Sigma-Aldrich) at 37 °C for 1 hour and then rinsed extensively. *Staphylococcus aureus* (ATCC25923) was cultivated overnight in lysogeny broth (LB) medium (10 g/l tryptone, 5 g/l yeast extract, 10 g/l sodium chloride, pH 7) at 37 °C. Bacterial cells were reseeded in LB medium and grown at 37 °C to reach the exponential growth phase (OD_600_ of 0.6 −0.7) followed by staining with CellTracker^™^ Green for 30 min and inactivation with 50% ethanol for 4 hours at 37 °C. Both pathogen stocks were stored at −20 °C until use.

#### *Ex vivo* whole-blood infection assay

Peripheral blood from healthy volunteers and from patients undergoing cardiac surgery with extracorporeal circulation was collected in S-monovettes^®^ (Sarstedt) containing recombinant Hirudin as anti-coagulant. Differential blood cell counts were measured with an auto hematology analyzer (BC-5300, Mindray). For whole-blood infection, PBS as mock-infected control (mock) or 1 × 10^6^ killed pathogens/ml whole blood were incubated for various time points (as indicated) at 37 °C on a rolling mixer (5 rpm). After incubation, the samples were immediately placed on ice. To collect plasma samples, whole-blood aliquots were centrifuged for 10 min at 10.000xg and 4 °C, and the resulting plasma was stored at −20 °C until further analysis.

#### Flow cytometry

Differential staining and flow cytometry was applied to identify distinct immune cell populations as well as to measure activation of immune cells and their association to pathogens. For surface antigen staining on the different immune cells, whole blood was stained with mouse anti-human CD3 (clone SK7, T cells), CD19 (clone HIB19, B cells), CD56 (clone MEM-188, NK cells), CD66b (clone G10F5, neutrophils), obtained from BioLegend, and CD69 (clone L78, early activation antigen, BD Biosciences). Monocytes were labelled with anti-human CD14 antibody (clone REA599, Miltenyi Biotec). Immature phenotype of neutrophils was assessed by surface CD10, CD16 and CD62L expression using mouse anti-human CD10 (clone HI10a, BioLegend), CD16 (clone 3G8, BioLegend) and CD62L (clone REA615, Miltenyi Biotec) antibodies. Surgery-induced changes on monocyte surface antigen expression were analysed for HLA-DR (anti-human HLA-DR, clone REA805, Miltenyi Biotec) and CD62L. Red blood cells were lysed with BD FACS Lysing solution followed by washing and harvesting cells in BD CellWASH solution. Acquisition was performed with the BD FACSCanto II flow cytometer and data was processed with FlowJo 7.6.4 software. The strategy used to evaluate the association of microorganisms to immune cells in human blood was shown for GFP-expressing *C. albicans* in Hünniger *et al*., 2014 and used in the same way for CellTracker^™^ Green-labeled *C. albicans* and *S. aureus* in this study.

#### Quantification of cytokines

The secretion of cytokines was assessed in plasma samples using Luminex technology (ProcartaPlex^™^ Multiplex Immunoassay, Thermo Fisher Scientific). The analyses were performed according to the instructions from the manufacturer.

### Statistical analyses

Data are presented as arithmetic means ± standard deviation (SD). Statistical analysis was performed by applying the following steps. First, the Shapiro-Wilk test was applied to test whether the underlying data is normally distributed. For normally distributed data the unpaired *t*-test was used to test for significant differences, since the respective data was unpaired. If the data was not normally distributed, either the Wicoxon signed-rank test was applied to test paired samples, or the Mann-Whitney U test was applied to test unpaired samples for significant differences. Afterwards, a multiple comparison correction (Bonferroni’s correction) was performed if comparisons were made between several data sets. The corrected *P*-value is given by P’. Significance is shown as \**P* < 0.05, \*\**P* < 0.01, \*\*\**P* < 0.001.

### Biomathematical Modeling

#### State-based virtual infection modeling

The state-based model (SBM) is derived from our previous models of whole-blood infection [16–18] and simulates the immune defence in whole blood that was infected with killed pathogens. As depicted in S4A Fig, the SBM contains several states for distinct cell populations in the system. Killed pathogens in extracellular space are represented by the state *P*_*KE*_ and pathogens that acquired immune escape are described by the state *P*_*IE*_. Furthermore, the model contains the states *N*_*j*_ and *M*_*j*_ that represent neutrophils and monocytes, respectively, where the index *j* denotes the number of phagocytosed pathogens. In order to compare the dynamics of the model with experimental data, we defined combined units that are specific combinations of measurable model states. Pathogens in extracellular space are represented by the combined unit *P*_*E*_:

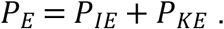

The combined units *P*_*N*_ and *P*_*M*_ refer to pathogens that have been phagocytosed by either neutrophils or monocytes:

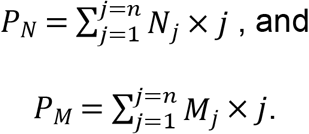

The transitions between model states represent biological processes during whole-blood infection (see the connections between states in S4A Fig). In the SBM, we defined three transition rates characterising specific transitions. The rate *φ*_*N*_ refers to the phagocytosis of killed pathogens (*P*_*KE*_) by neutrophils with *j* phagocytosed pathogens (*N*_*j*_) and characterises the state transition

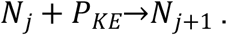

The phagocytosis of killed pathogens (*P*_*KE*_) by monocytes with *j* phagocytosed pathogens (*M*_*j*_) is characterised by the rate *φ*_*M*_ and is described by

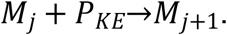

The immune escape of killed pathogens (*P*_*KE*_) is quantified by the rate *ρ* for state transition

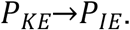

The transition rates enables us to quantify the dynamics of the SBM, where a transition from any state *S* to state *S′* within the time step Δ*t* will be performed with probability *P*_*S*→*S*′_: *P*_*S*→*S*′_= *r*_*S*→*S*′_×Δ*t*. The SBM dynamics was calculated by applying the random selection method [46] and the flow chart of this simulation algorithm is depicted in S5A Fig.

In order to estimate the *a priori* unknown values of transition rates, the state-based model (SBM) is fitted to experimental data of the whole-blood infection assays by applying the algorithm of *simulated annealing* [47] based on the *Metropolis Monte Carlo* scheme [48]. For a detailed description of the parameter estimation in the SBM we refer to the Supplementary Information and to S6A Fig.

### Agent-based virtual infection modeling

In order to investigate also spatial aspects of host-pathogen interactions we applied a previously developed agent-based model (ABM) [49,50] that was adjusted to the context of whole-blood infection assays [17,18]. This spatial counterpart of the SBM simulates the different immune cell types of neutrophils and monocytes as well as the killed pathogens as distinct spherical objects, i.e. the agents (S4B Fig). These agents can migrate and interact within an environment in a rule-based fashion, where the environment is a continuous three-dimensional representation of a section of 0.5 μ*l* of the whole-blood infection assay in which cells perform random walk migration due to the high density of erythrocytes [17]. Modeling the active migration of neutrophils and monocytes by a diffusion process with coefficients *D*_*N*_ and *D*_*M*_, respectively, the passive migration of pathogens is estimated to have a small diffusion coefficient of *D*_*P*_ ≈ 1 μ*m*^2^/*min* [17]. Based on the random selection method [46], each stochastic simulation was repeated 30 times to gain statistically sound results (S5B Fig). The migration parameters of immune cells were estimated from a minimization of least-squares error (LSE) between experiment and simulation using the method of *adaptive regular grid search* [51] (see Supporting Information and S6B Fig).

## Acknowledgments

We are grateful to Prof. Torsten Doenst from the Department of Cardiothoracic Surgery for his support of the project and to Petra Bloos, Anja Haucke, Karina Knuhr-Kohlberg, Steffi Kolanos and Katrin Schwope for organizing blood draws on the ICU. We thank Carl-Magnus Svensson for scientific discussions throughout the project.

---

**S1 Figure.**
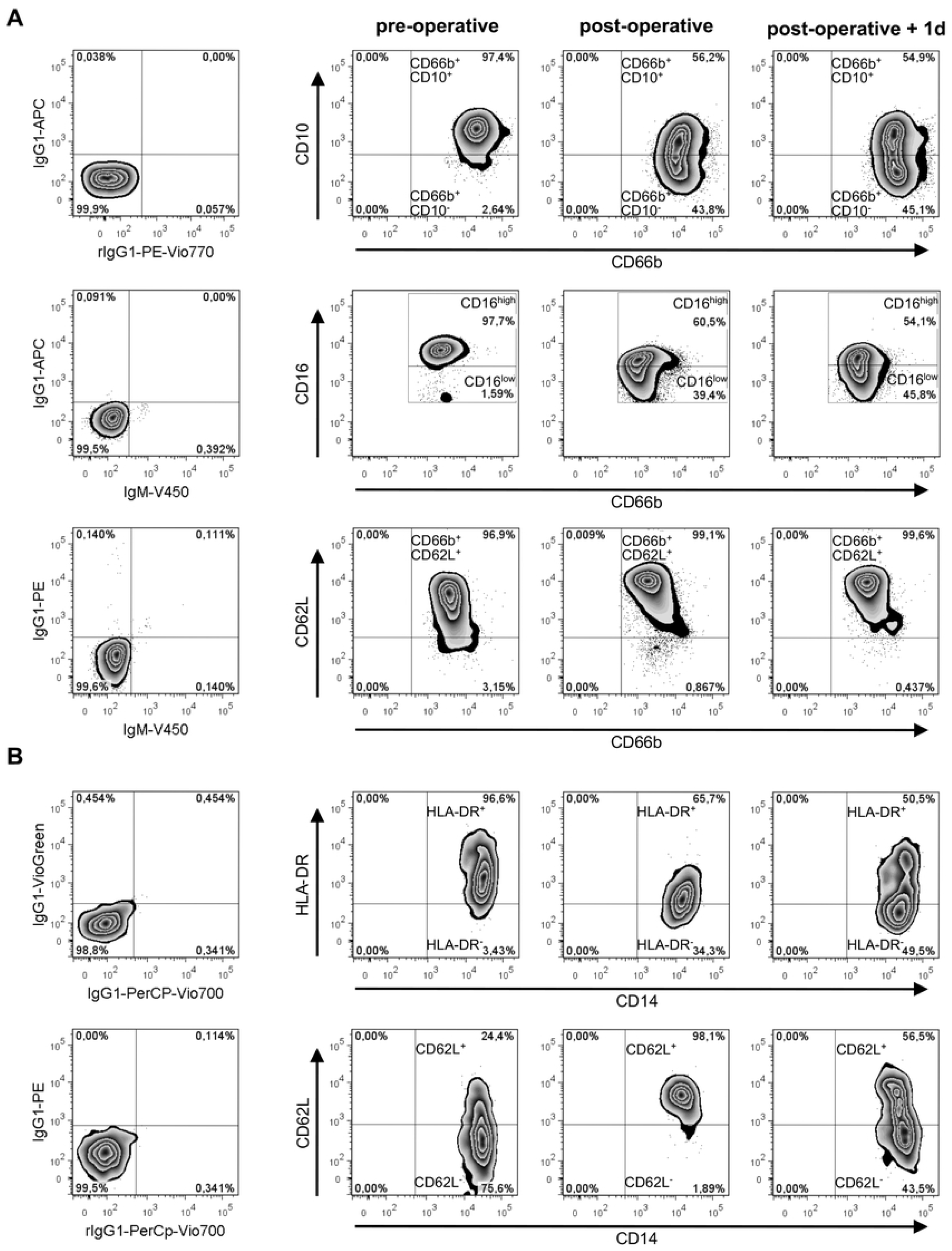

**S2 Figure.**
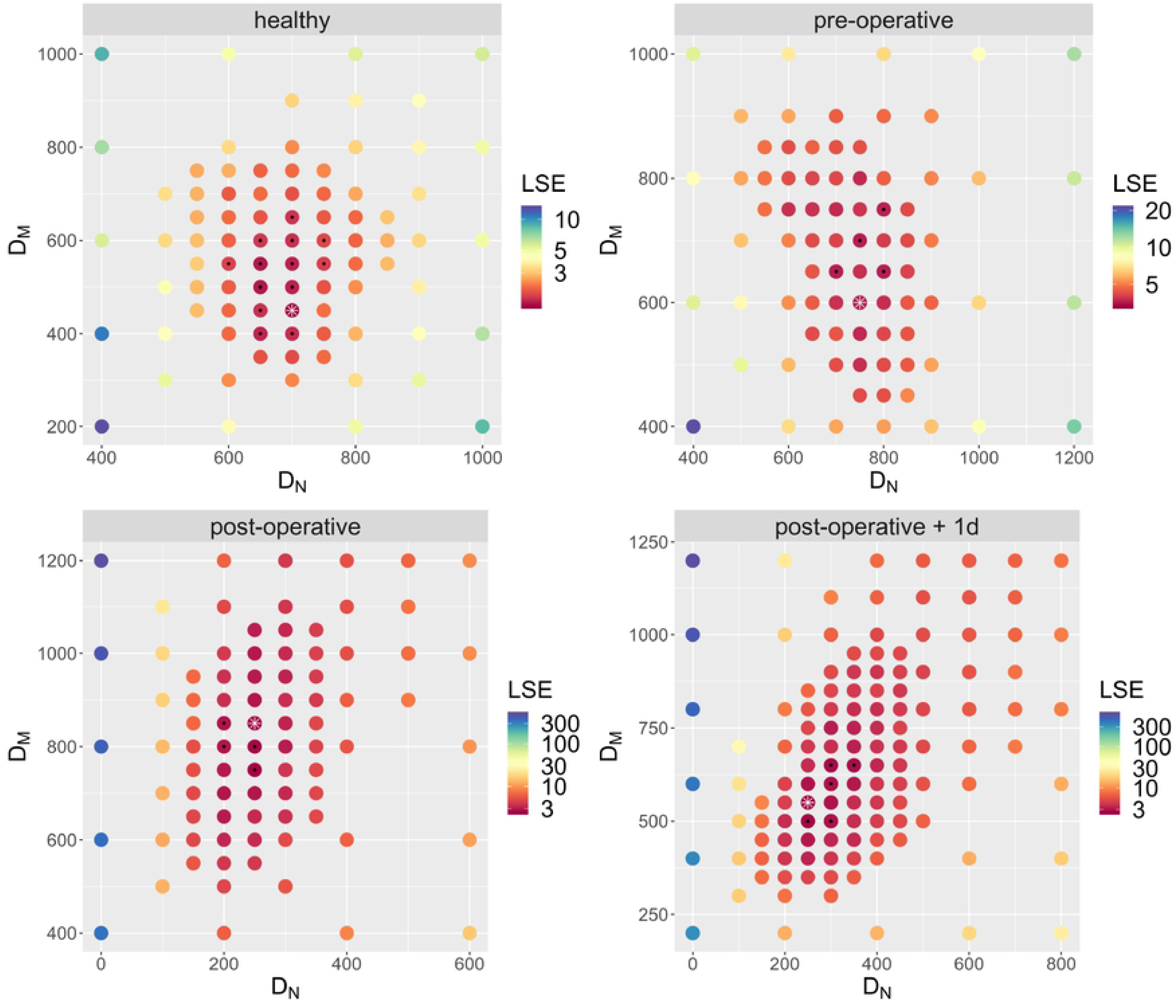

**S3 Figure.**
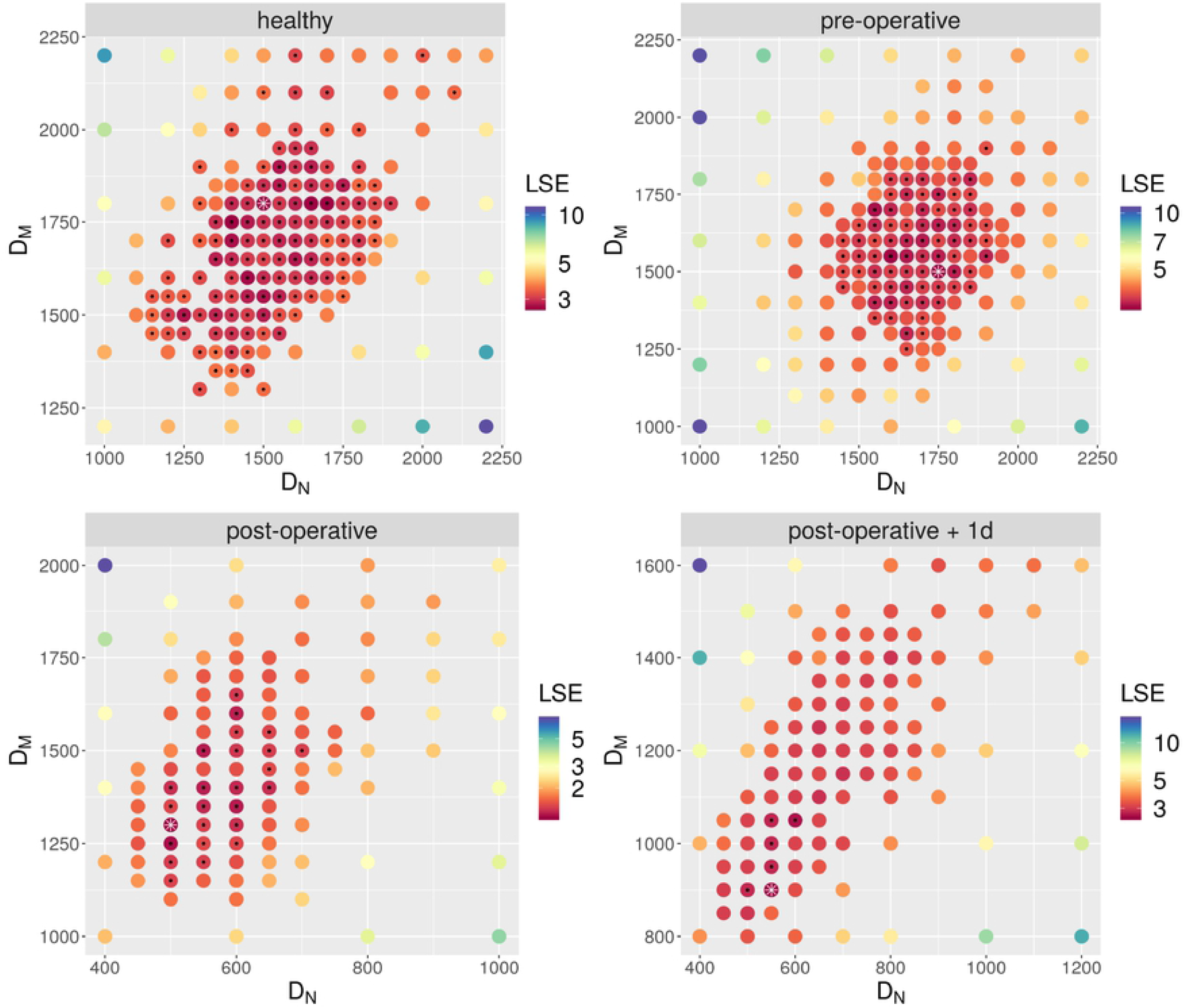

**S4 Figure.**
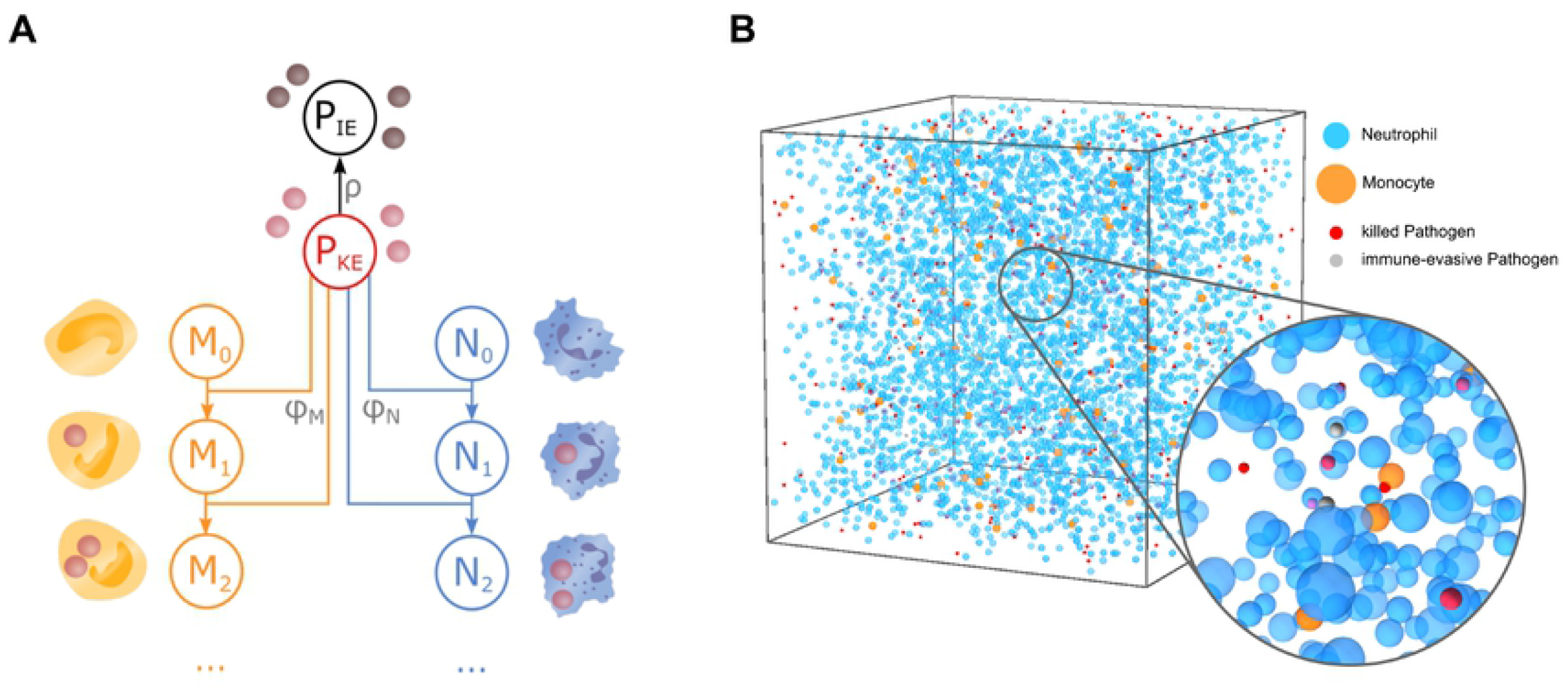

**S5 Figure.**
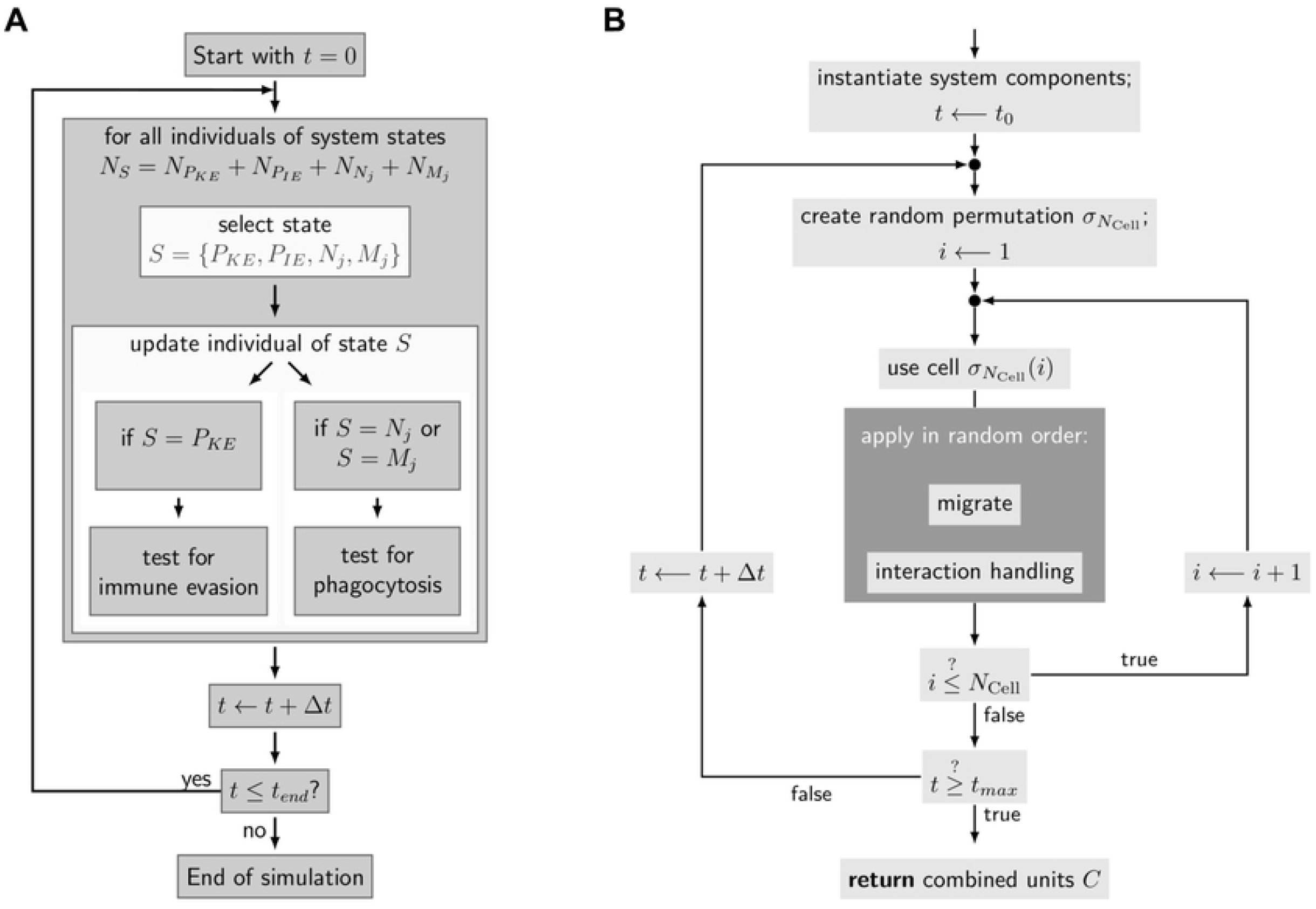

**S6 Figure.**
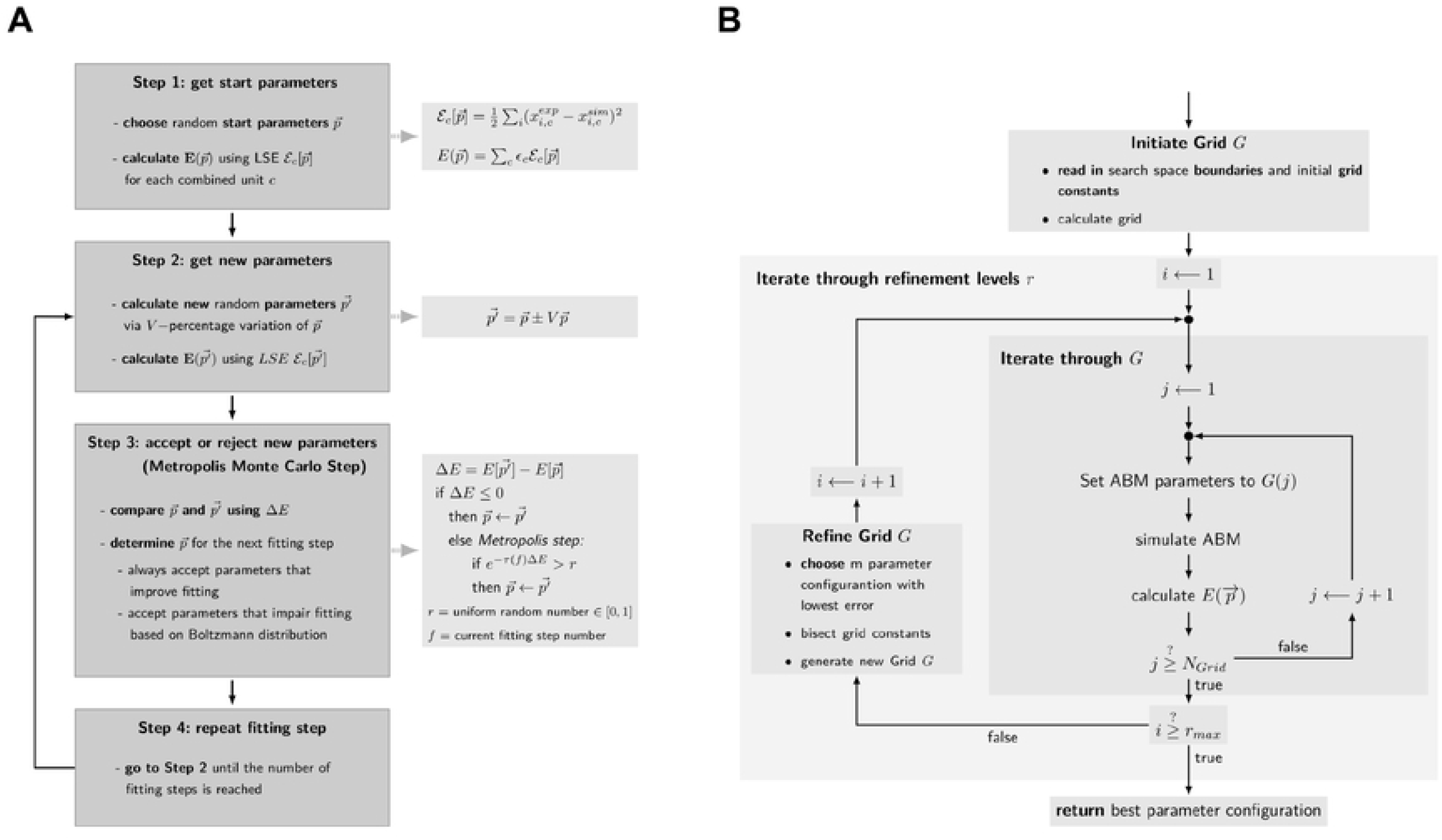

